# Brendo: an open resource for uniquely transcribed genes in brain endothelial cells

**DOI:** 10.1101/2025.10.25.684508

**Authors:** Ozgur Beker, Fereshteh Ramezani Khorsand, Ogun Adebali, Nur Mustafaoglu

## Abstract

The blood-brain barrier (BBB) plays a critical role in central nervous system homeostasis, yet comprehensive transcriptional profiling of brain microvascular endothelial cells (BMECs) remains limited. Leveraging the increasing availability of publicly accessible bulk RNA sequencing (RNA-seq) datasets, we developed an integrated analytical framework to identify genes selectively enriched in BMECs compared to endothelial cells (ECs) from other tissues. To address the substantial batch effects inherent to multi-source bulk RNA-seq data, we combined two differential expression strategies, rank aggregation methods, and extensive quality controls. Our analysis incorporated EC samples from 16 different tissues and employed robust statistical workflows to mitigate technical confounders while preserving biologically meaningful signals. Validation using known BBB markers and independent proteomic evidence confirmed the reliability of our approach.

We present Brendo (Brain Endothelial Open Resource), an open-access web platform providing searchable, filterable, and downloadable data on differentially expressed genes in BMECs. Brendo enables an in-depth exploration of brain endothelial gene expression and offers broader applications across vascular biology by supporting cross-tissue EC comparisons. The workflow described is adaptable to other biological contexts, promoting the systematic reuse of public bulk RNA-seq datasets for target discovery. Collectively, this resource provides a foundation for advancing BBB-targeted therapeutic strategies and highlights the value of open data integration in transcriptomic research.

## Introduction

Effective treatment of brain disorders requires the delivery of therapeutic agents across the systemic circulation and, in particular, successful traversal of the blood-brain barrier (BBB). The BBB is a specialized form of vasculature in the central nervous system (CNS) that establishes chemical, physical, biological, and immunological barriers between blood and the brain [1, 2]. While this barrier protects the CNS from harmful substances, it poses a significant challenge for drug delivery due to its highly selective permeability. Anatomically, the BBB consists of several specialized cellular components, including brain microvascular endothelial cells (BMECs), which are tightly regulated by surrounding pericytes that contribute to vascular stability, and astrocytic end-feet that mediate interactions with neurons [1, 3–5]. Among these, BMECs play a pivotal role in maintaining BBB integrity by actively restricting the entry of most circulating molecules into the brain parenchyma [6–8]. To support this unique function, BMECs exhibit distinct molecular and structural features compared to endothelial cells in other organs. They form exceptionally tight junctions through the expression of specialized proteins such as claudins, occludins, and zonula occludens, along with adherens junction proteins, to prevent paracellular leakage [8–12]. Additionally, BMECs express various efflux transporters at high levels, contributing to their restrictive nature [13–15]. These features are reflected in unique RNA and protein expression profiles as well as their functional assays that differentiate BMECs from other endothelial populations.

Overcoming the BBB remains one of the greatest challenges in the treatment of CNS diseases. Although many innovative strategies are being explored – both invasive and noninvasive – noninvasive approaches are generally preferred due to their safety and clinical feasibility. However, no noninvasive method has yet received clinical approval. Among these strategies, receptor-mediated transport (RMT) represents a particularly promising approach for delivering therapeutics across the BBB [16–18]. RMT leverages endogenous receptors, such as the insulin receptor (IR) and the transferrin receptor (TfR), to enable the targeted transport of drugs into the brain [19, 20] . Although several RMT-associated proteins – including IR and TfR – have been extensively studied by academic and pharmaceutical researchers, one of the primary limitations is their widespread expression across multiple cell types and tissues, which reduced their specificity to BMECs [21, 22]. Therefore, the success of RMT-based drug delivery depends on the identification of genes that are preferentially expressed in BMECs. These genes offer targets that can enhance specificity, minimize off-target effects, and accelerate the development of CNS therapeutics. Thus, a tool that identifies differential gene expression between BMECs and other endothelial cell types, taking into account their subcellular localization and transport potential, is needed to identify promising targets for shuttling drugs and/or drug carriers across the BBB. In this context, bulk RNA-sequencing (RNA-Seq) has emerged as a powerful tool to investigate gene expression profiles across tissues and cell types.

In this study, we curated a dataset comprising 264 publicly available, healthy human bulk RNA-Seq samples from 63 different studies. These samples represent endothelial cells from 16 tissues (Figure 1). We performed pairwise differential expression analyses between BMECs and each of the 15 other endothelial tissues using both DESeq2 [23] and the Wilcoxon rank-sum test (WRST). We then aggregated the results to determine the number of tissues each gene showed BMEC-specific over-or under-expression, along with average expression values. To ensure robustness, we performed bootstrapping by randomly selecting %60 of BMEC samples and repeating the full analysis 100 times. Aggregated results from these bootstraps were used to assess the consistency of differential expression. To refine our focus for identifying targeting proteins that have the ability to cross the BBB, we prioritized genes encoding transmembrane proteins, which are especially relevant for RMT mechanisms [17, 24, 25].

**Figure 1.**
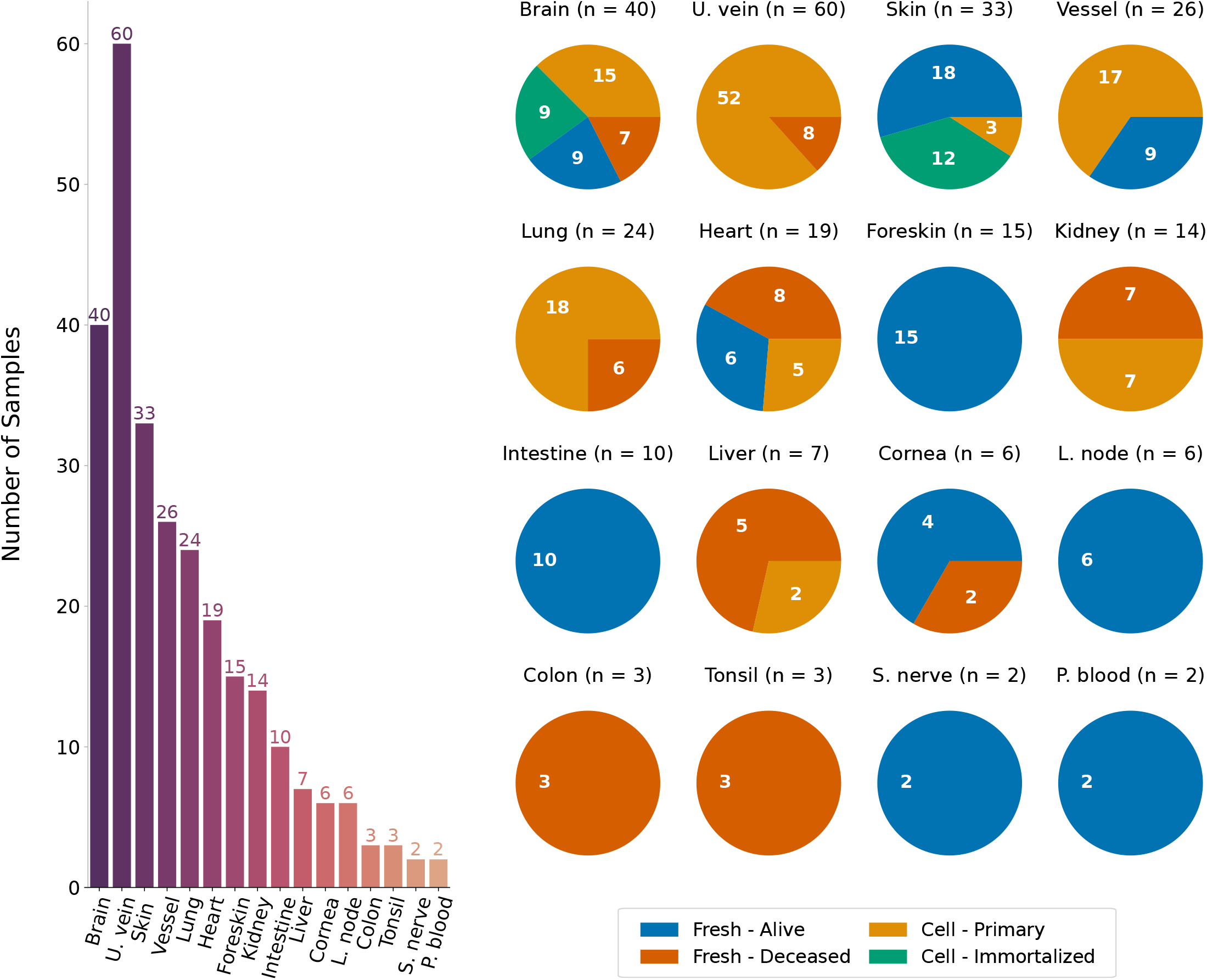
Distribution of samples by tissue and cell source. Only samples retained throughout the entire workflow are represented. Left: Number of samples across all tissues included in the study. Right: Distribution of samples across four distinct cell sources: fresh tissue from living individuals, fresh tissue from deceased individuals, primary cell lines, and immortalized cell lines.

Based on these analyses, we present Brendo—an open-access resource for cataloging genes uniquely transcribed in brain endothelial cells. We envision Brendo as a foundational tool to accelerate the development of BBB-targeted therapeutic strategies. Moreover, Brendo serves as a versatile platform for comparing endothelial gene expression across 15 other tissues, potentially advancing drug delivery strategies beyond the brain and into other organs.

## Results

### Data Curation

We started by curating a dataset of 264 healthy endothelial cell samples from 63 studies focusing on different tissues (*Methods*). Figure 1 describes the resulting distribution of samples across tissues (left) and cell sources (right) for the final dataset. Median-ratio normalized (MRN) counts across samples are visualized in Figure 2. We ran a separate analysis for a subset of 140 samples – originating from the BBB, colon, heart, intestine, kidney, liver, lung and vessel tissues – for increased focus. MRN normalization was performed separately for this analysis, using only the considered samples.

**Figure 2.**
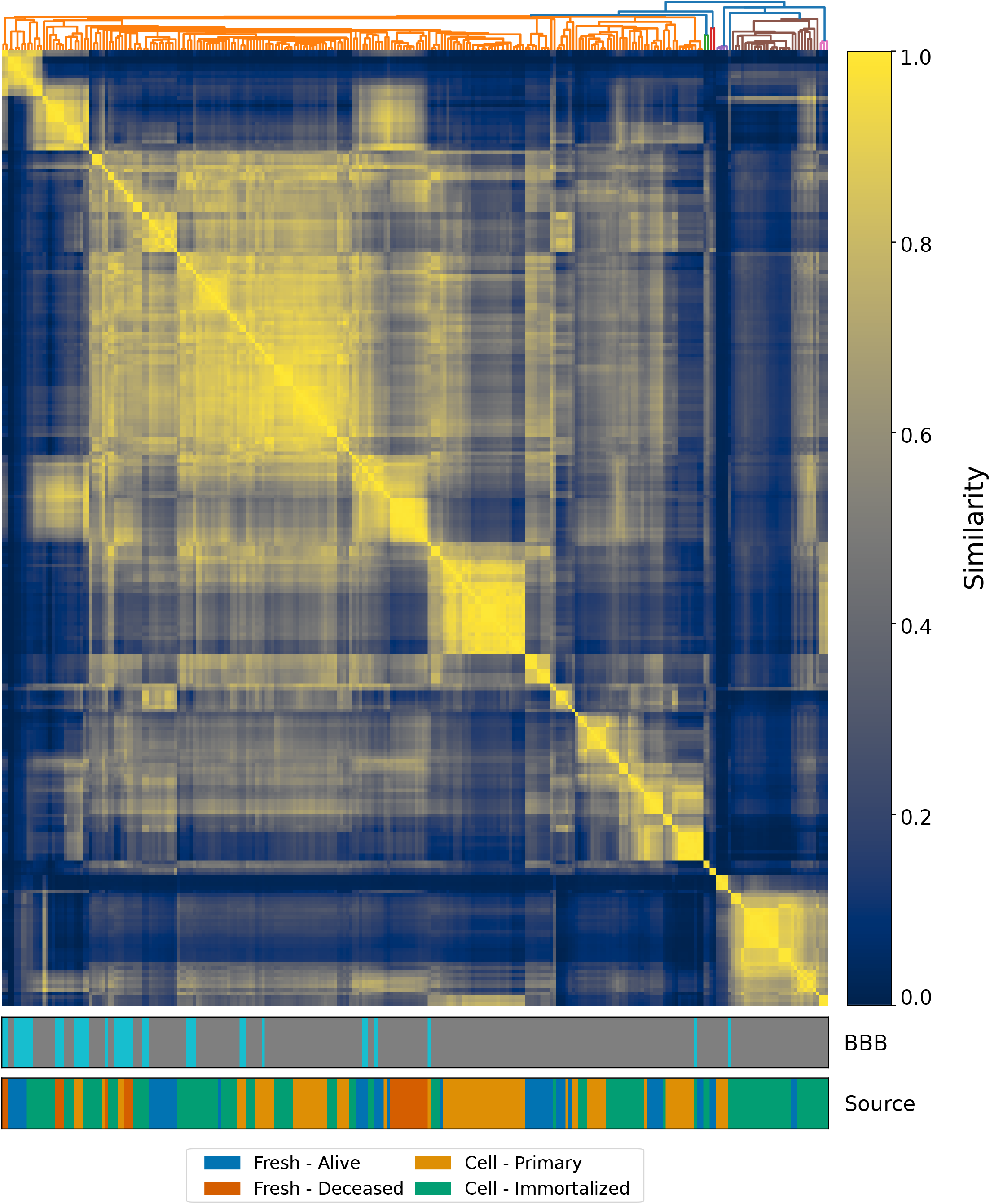
Sample relationships for the curated dataset. Samples were clustered using single linkage based on pairwise cosine similarities of MRN normalized counts. Rows and columns represent individual samples. Bottom color bars indicate BBB-derived samples (BBB) and sample source (Source).

### Statistical Analysis

#### Differential Expression Testing

We employed two complementary statistical methods — DESeq2 (v1.34) [23] and the WRST — to conduct statistical testing for differential gene expression in pairwise comparisons between brain and other tissues (*Methods*). This decision was motivated by previous permutation analyses demonstrating a high rate of false positives in population-level datasets [26]. We present the results in Figure 3. Here, each approach is a combination of the testing method used and the set of tissues considered. Notably, DESeq2 consistently identifies more genes as differentially expressed for the same tissue set, whereas WRST did not identify any gene as significant across more than eight comparisons.

**Figure 3.**
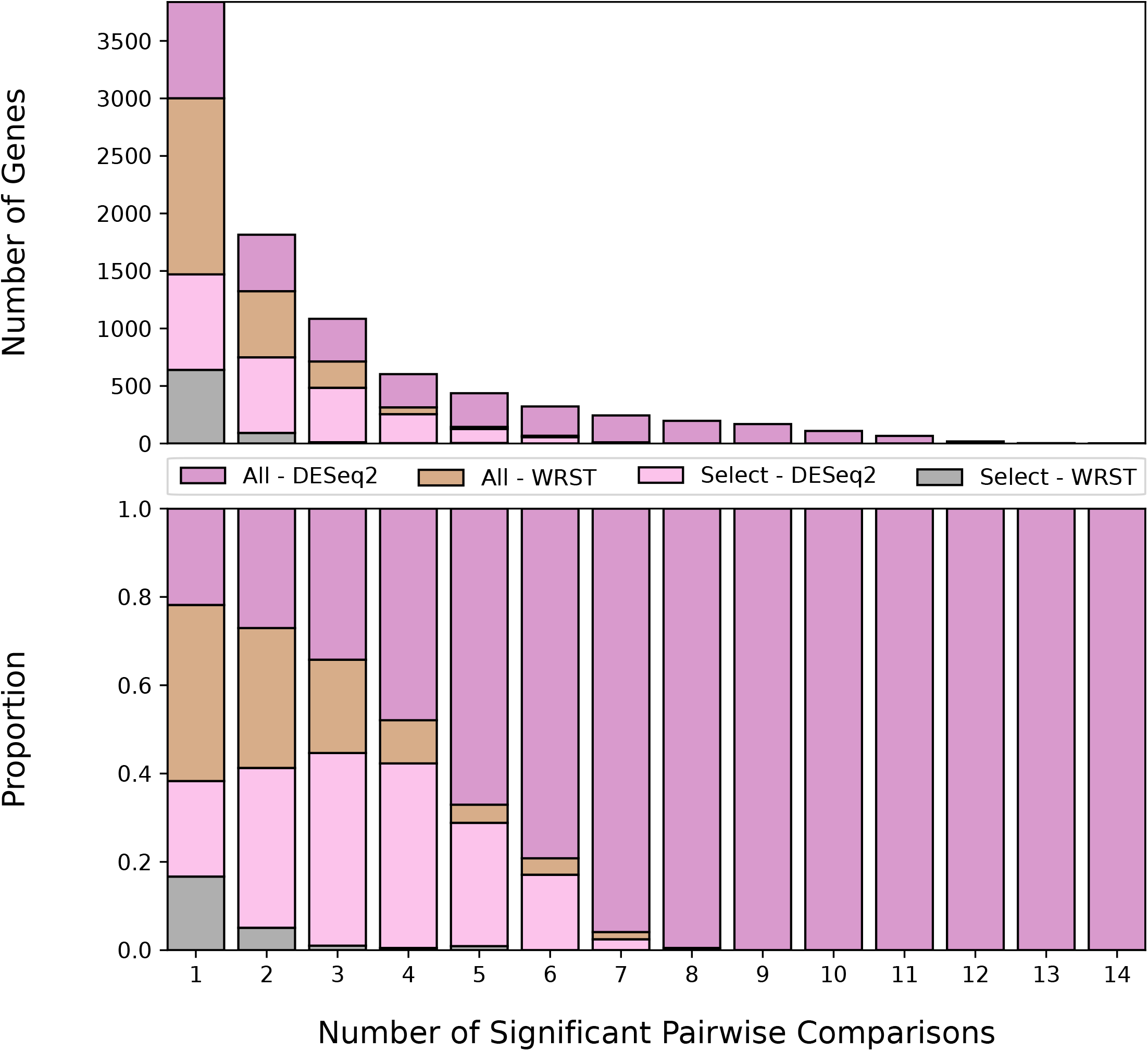
Differentially overexpressed genes in the BBB. Bar plots indicating the number of significantly enriched genes (top) and its proportion across all approaches (bottom) for each approach.

#### Aggregating Differential Expression Results

To consolidate our findings across method outputs, we applied Robust Rank Aggregation (RRA, *Methods*) [27]. Primary aggregation focused on four merged lists corresponding to over-expressed transmembrane-helix-containing genes, stratified by the statistical methods and dataset scope. Two additional aggregations were performed for each statistical method separately to further prioritize candidates. To validate the robustness of our findings, we rerun the analysis by bootstrapping the original BMEC samples (*Methods*). Top 10 significantly enriched genes from RRA and bootstrap validation results are presented in Figure 4, integrated into the Brendo platform.

**Figure 4.**
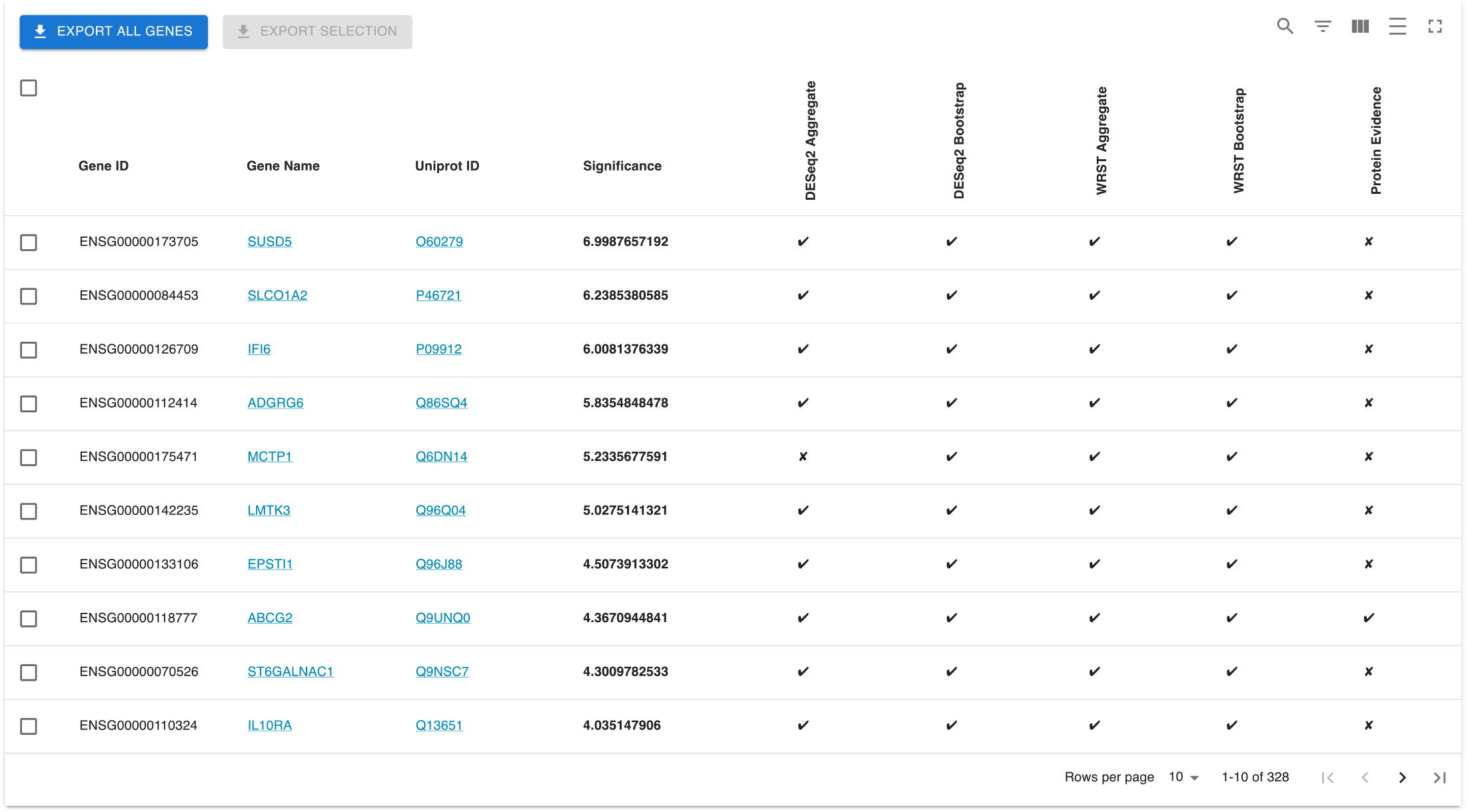
Screenshot of the RRA and bootstrap results from Brendo RRA and bootstrap validation results are listed for top 10 transmembrane helix filtered genes. Additional results are available at the platform webpage.

#### Gene Ontology (GO) Enrichment Analysis

We performed GO enrichment analysis (*Methods*) on the prioritized gene list using PANTHER [28] and its GO enrichment testing tool [29]. Figure 5 depicts the GO enrichment results for the final prioritized gene list. Notably, we observed the ontology “transport across the BBB” being significantly enriched, pointing to the validity of results obtained by our analysis.

**Figure 5.**
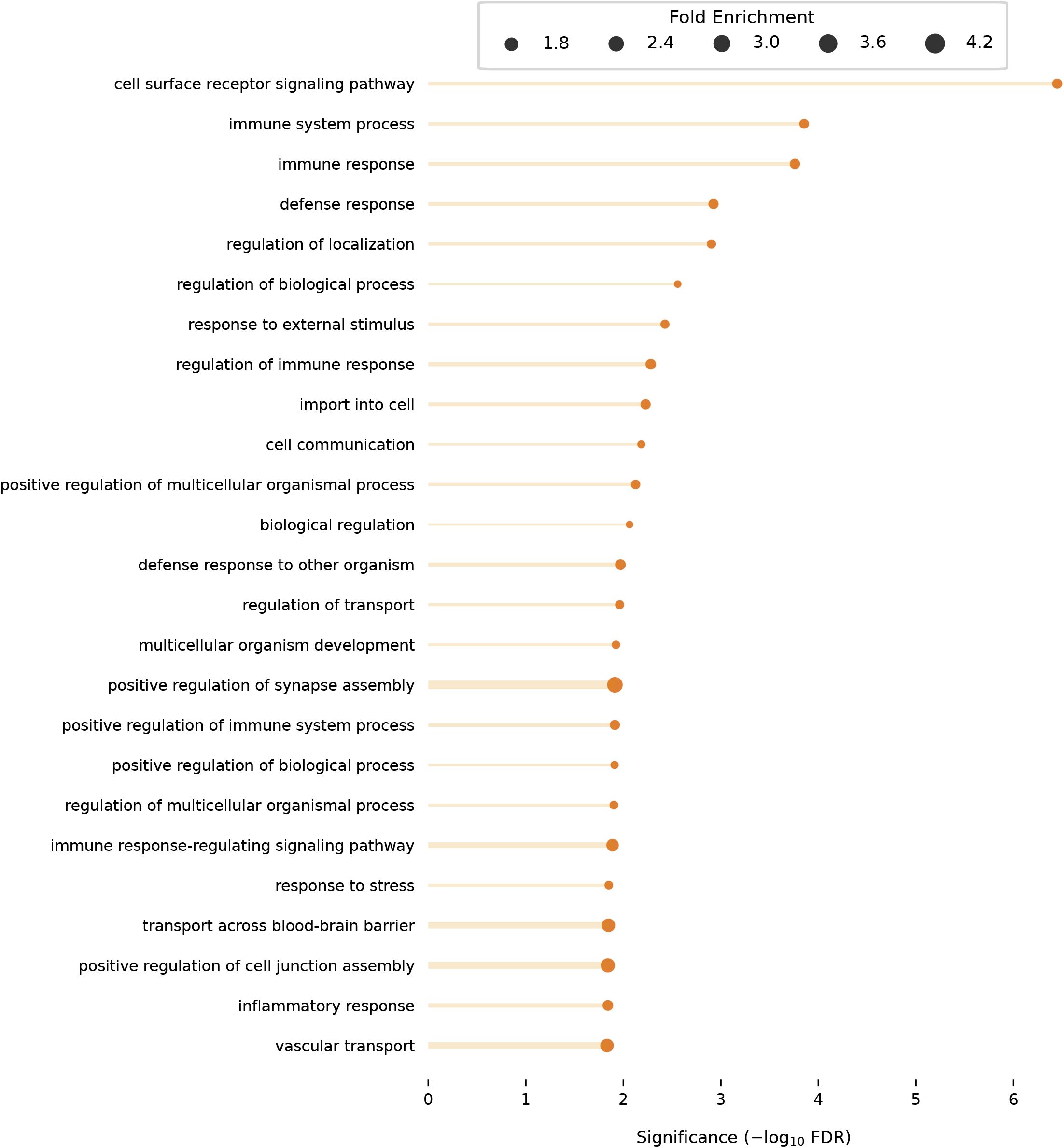
Gene ontology (GO) enrichment analysis of prioritized BBB genes. The top 25 significantly enriched terms (− log_10_ FDR *<* 0.05) are shown for genes retained after RRA and transmembrane helix filtering. Entries are sorted by significance (− log_10_ FDR). Circle size and bar thickness both represent fold enrichment.

### Brendo Platform

#### Technical Details

The Brendo platform operates on a FastAPI [30] back end with a ReactJS [31] front end. It can be reached online at brendo.sabanciuniv.edu, ran locally or be deployed on proprietary servers by following the instructions provided in the repository. For specific requests or detailed inquires regarding implementation, users are encouraged to consult the GitHub page (Code and Data Availability).

#### Platform Structure

The platform is organized into several tabs. The “Transmembrane Focus” presents extended results obtained through the previously described methods, specifically for genes encoding proteins with predicted trans-membrane helices that are overexpressed in BMEC samples. This table includes information columns from: (1) our statistical analysis, (2) independent protein evidence through the public PXD018602 dataset and (3) a perfusion score obtained by aggregating blood perfusion rates for certain organs (Methods). The “All Genes” tab contains merged results across all combinations of three binary conditions (overexpression/underexpression, full dataset/selected tissues, statistical test used), excluding transmembrane filtering.

#### Filter and Search Capabilites

“Transmembrane Focus” and “All Genes” tabs support extensive user interactions, including:

- String-based search
- Factor-based filtering (“present”, “absent” or “do not care”)
- Sorting for numeric columns
- Exporting results for selected (or all) genes.
- Quick links to search a gene name in the Human Protein Atlas [32] or a Uniprot ID in InterPro [33]
- MRN-normalized counts across all tissues by clicking on significance or fold change values for a specific gene, presented in Figure 6.

**Figure 6.**
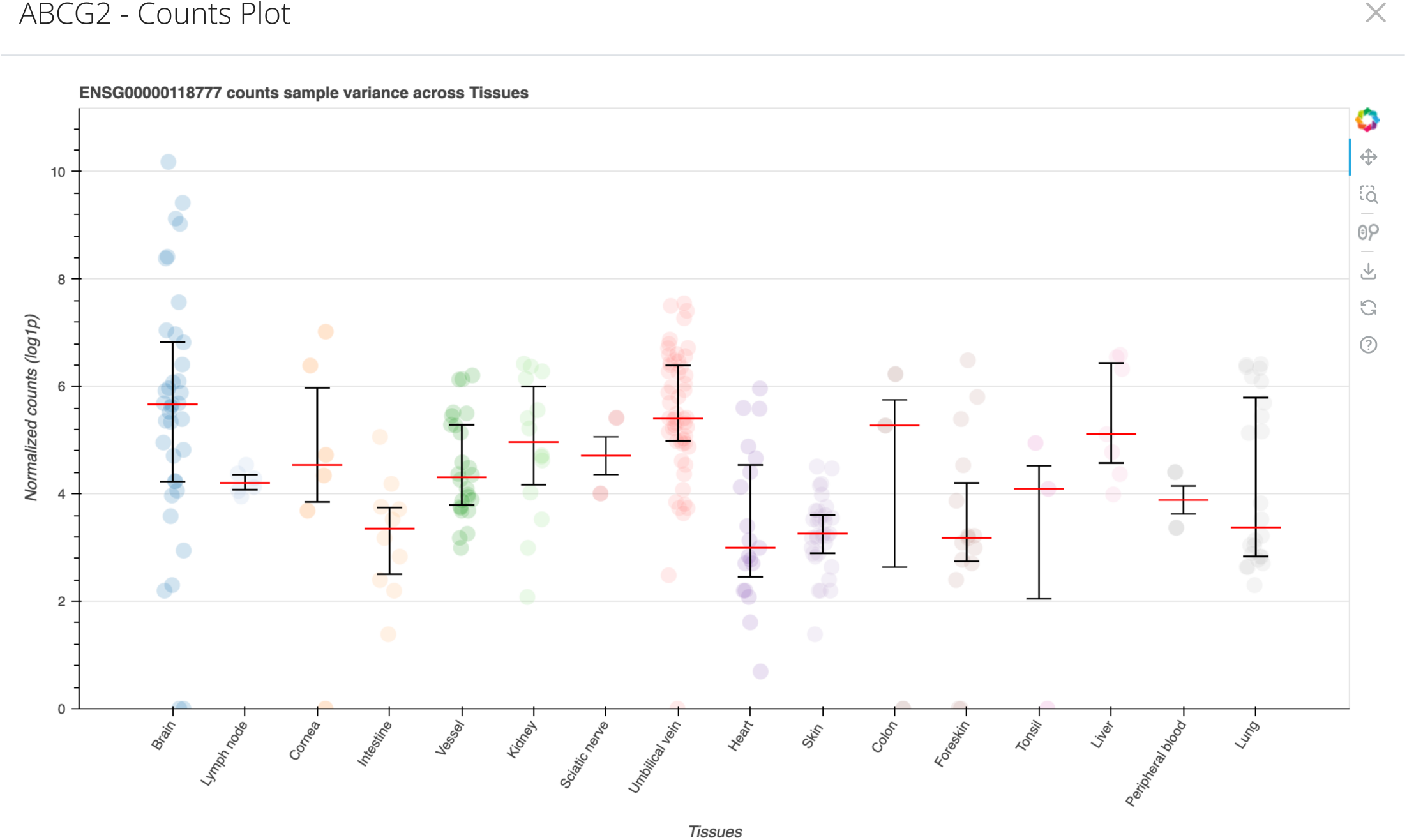
Screenshot of the count distribution plot display for ABCG2 (ENSG00000118777). Users can access MRN-normalized counts and visualize their distribution across tissues for genes displayed in a table by clicking on their significance or fold-change values.

We provide a summary of the our Methods through the “Methods” tab. Users can also consult the “Support” tab for contact information, citation guidelines, and future announcements.

## Discussion

The increasing availability of publicly accessible RNA-Seq datasets has enabled comprehensive analyses of baseline and differential expression. Resources such as the Expression Atlas [34] provide broad expression profiles across various tissues and experimental conditions, facilitating candidate gene identification. While more specialized databases—such as those focused on brain or BBB-specific expression— are available, resources like BBBomics [35] often suffer from limited sample sizes and inadequate representation of cell-type diversity. A major challenge in large-scale transcriptomic analyses is the presence of batch effects—systematic technical variations unrelated to biological differences. These confounders, arising from differences in library preparation, sequencing platforms, or laboratory protocols, can significantly bias results. Therefore, it is essential to apply robust statistical methods to quantify and validate differential expression while mitigating technical variation.

In our analyses, we confirmed that DESeq2 consistently identified a larger number of significantly differentially expressed genes and yielded lower adjusted p-values compared to WRST, particularly when the model intercept was left unconstrained. Genes previously confirmed to be expressed in the brain at RNA and protein levels, such as ABCG2, ABCB1, and SLC2A1 [7, 36, 37] were consistently identified and ranked higher by both DESeq2 and WRST. However, notable discrepancies in gene rankings were observed between the two methods, with WRST yielding fewer significant genes overall.

Restricting the scope of tissues under comparison influenced MRN-normalized read counts and reduced the number of significant comparisons per gene but had minimal impact on final gene rankings. RRA emerged as a reliable method for consolidating rankings across varied conditions. Notably, top-ranking genes in the final list of transmembrane-helix-containing candidates were predominantly supported by prior literature or protein evidence from PXD018602 BMEC samples.

A particular strength of this study lies in its integration of baseline healthy samples from diverse sources, aiming to capture natural population variance. However, this approach also amplifies batch effects, as described earlier. To address this, we enforced a zero-intercept model in DESeq2, which yielded results more consistent with WRST, reduced the number of false-positive findings, and minimized biases associated with technical variation.

During pairwise comparisons, we observed overexpression of epithelial markers in iBMEC samples relative to primary BMECs, previously pointed out by Lu et al. [38]. Since differentiation of BMECs from iPSCs is a challenging task and the protocols may need to be further improved, the outcomes of some of the differentiation protocols may lead to cells with epithelial characteristics, and our differential analyses between iBMECs and other BMECs corroborated earlier findings [38, 39] regarding potential misidentification of some iBMEC samples. To avoid introducing artifacts into our results, we excluded iBMECs from all subsequent analyses.

Finally, although this study focuses on ECs, the analytical workflow is broadly adaptable. By adjusting the reference group and selecting appropriate datasets, the approach can be generalized to identify differential expression patterns in other tissues or cell types using publicly available bulk RNA-seq data.

## Limitations

We believe several limitations of our approach warrant discussion. First, in bulk RNA-seq samples, the proportion of ECs cannot be precisely quantified, raising the possibility of expression contamination from other brain-resident cell types. Second, uneven sample sizes across tissues complicate the interpretation of certain pairwise comparisons where BMECs were compared against much smaller sample groups. To mitigate these limitations, we cross-validated key findings through multiple strategies: incorporating protein expression evidence (e.g., for ABCG2 and SLC2A1 in BMECs), performing Gene Ontology (GO) enrichment analysis, and cross-referencing top-ranked genes with prior studies to ensure biological plausibility.

## Conclusion

Within this study, we sought to determine differential expression patterns in BMECs using publicly available bulk RNA-seq data from ECs across various tissues. Despite the challenges posed by batch effects and inaccuracies inherent to large-scale bulk RNA-seq analyses, we successfully identified expression patterns for both previously characterized and novel genes and provided supporting evidence for the robustness of our findings. All results have been made publicly available through the Brendo web service to serve as a reference for future research in the field. Additionally, we discussed the adaptability of the workflow for conducting similar analyses across different biological contexts. Consequently, this study highlights the value of open data practices and further encourages the adoption of similar approaches within the research community.

## Methods

### Data Curation

#### Raw Data

Initially, we collected a total of 321 healthy samples (after summing technical replicates) representing 16 tissues across 74 studies from the Sequence Read Archive (SRA) and Gene Expression Omnibus (GEO) databases. Sample sources included primary cell lines (CP), immortalized cell lines (CI), induced pluripotent stem-cell derived samples (CS), and fresh tissue extracted from deceased (FD) or live (FA) individuals. Fifteen of these samples contained paired-end, single-indexed reads generated through various single-cell sequencing platforms. Read files were downloaded through the European Nucleotide Archive (ENA) Fast Adaptive and Secure Protocol (FASP) server when available, and through the SRA Toolkit otherwise.

#### Filtering and Quality Control

Reads were aligned to Homo sapiens reference genome (GRCh38 - Release 107 primary assembly and annotations) using STAR v2.7.0 [40], with the `sjdbOverhang` parameter set to 100. Alignment quantification was performed in R [41] using the Rsubread v2.8.2 [42] function `featureCounts`. For downstream analyses, only samples with a minimum of 10,000 raw transcript counts across genes and genes with at least 5 total counts across samples were retained. BMEC samples derived from induced pluripotent stem cells (iPSCs) were excluded due to potential epithelial-like properties previously reported by Lu et al. [38].

#### Data Curation

In total, we established a final dataset of 264 healthy samples from 63 studies for subsequent analyses. Of these, 40 samples belong to the BMEC group. Other samples belonged to endothelial cells (ECs) from 15 distinct tissues – colonic microvascular ECs (colon), corneal ECs (cornea), cardiac and aortic valve ECs (heart), mucosal ECs (intestine), glomerular and peritubular ECs (kidney), sinusoidal ECs (liver), pulmonary arterial ECs (lung), lymphatic and microvascular ECs (foreskin), lymphatic ECs (lymph node), peripheral blood ECs, endoneurial ECs (sciatic nerve), dermal microvascular and lymphatic ECs (skin), tonsillar ECs, umbilical vein ECs, and arterial ECs (vessels) [36, 38, 43–92].

### Statistical Methods

#### DESeq2

DESeq2 (v1.34) [23] models gene counts by fitting a negative binomial generalized linear model (GLM) with a logarithmic link function. For our case, the design is not factorial but comparative, thus we set the intercept explicitly to 0. We fit a single model across all samples, and then extract pairwise contrasts post-fitting.

#### Wilcoxon rank-sum test (WRST)

Given ordinal and i.i.d. samples from two populations, the WRST provides a simple method for assessing whether a statistically significant difference exists between the populations. We conduct WRSTs using the approach outlined by Li et al. [26], treating samples from different tissue types as populations and MRN normalized counts as observations.

#### Merging Pairwise Comparisons

Method outputs were merged by calculating (1) the number of times each gene appeared significant across comparisons, (2) the mean fold change (log_2_), and (3) the mean significance (− log_10_ *padj*). Separate filtered result lists were also generated for genes encoding proteins with predicted transmembrane helices. In total, 16 merged results were produced, representing all combinations of four binary conditions: (1) statistical test used, dataset scope (full vs. selected tissues), (3) direction of differential expression (overexpressed vs underexpressed in brain samples), and (4) transmembrane helix filtering.

#### Robust Rank Aggregation (RRA)

Robust rank aggregation [27] is a method that provides statistical testing for genes that are ranked consistently higher across multiple ranked lists, under the null hypothesis that all ranked lists are randomly generated. We consider genes significant for a single run of RRA if they achieve *p <* 0.05.

#### Bootstrap Validation

We randomly downsampled %60 of BMEC samples without replacement 100 times and repeated the entire analysis pipeline on selected tissues, generating 200 additional merged results (100 per statistical method). Like before, we aggregate bootstrap results using RRA. Genes were significantly enriched for the validation step if they achieved an RRA *p <* 0.05.

#### GO Enrichment Analysis

GO enrichment analysis was conducted using the enrichment analysis tool available at PANTHER [28, 29]. For our purposes, we focus on the subcategory “biological process (BP)” for enrichment testing. GO results were considered enriched if they attained − log_10_ FDR *<* 0.05.

### Additional Information for Brendo

#### Protein Evidence

We derived the “Protein Evidence” column from detected proteins using label-free proteomic data of fresh human BMEC samples [93] processed using MaxQuant (v2.3.0.0) [94], with configuration details available in the Proteomics Identification Database (PRIDE) under the accession PXD018602.

#### Perfusion Scores

We calculated perfusion score for a single gene by summing the perfusion rates of its significant appearances across liver, heart, kidney and lung comparisons. Perfusion rates were derived by combining tissue density and volume information [95] with blood flow measurements [96], reported in units of 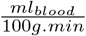.

## Code and Data Availability

The Brendo platform is freely available under the domain brendo.sabanciuniv.edu, while the code for [data processing scripts and the website source code](https://github.com/CompGenomeLab/brendo) are made available over GitHub.

Raw read & count data can be obtained by using the information from Table 1, and can be provided on request along with the metadata table used for the study.

**Table 1.**
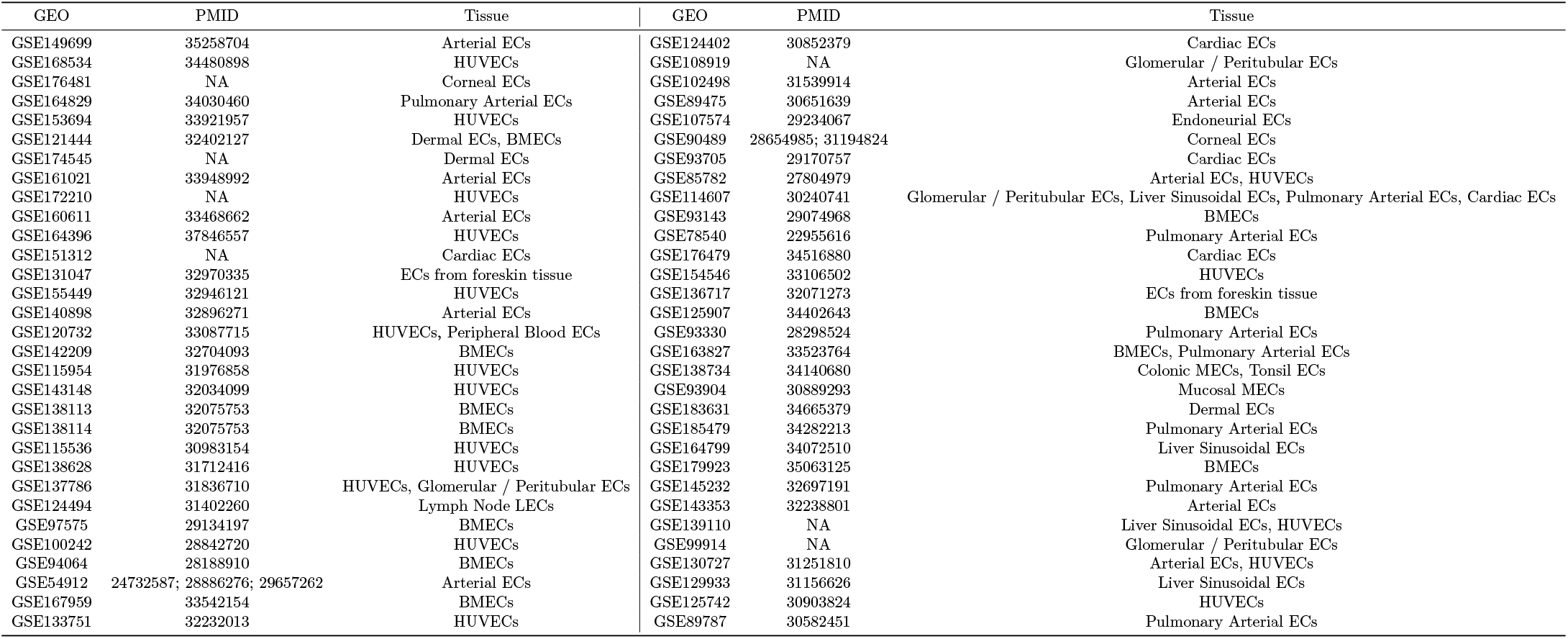
Metadata for all samples included initially in the dataset. This list includes all samples initially included in the study, but not all of the samples are retained for downstream analyses as underlined in Methods - Data.

## Acknowledgements

We want to thank Tandaç Furkan Güçlü and Zeynep Kılınç for their valuable inputs throughout the study. All computations were performed using the TOSUN cluster at Sabanci University.

## Funding

This research was supported by funding from Scientific and Technological Research Council of Türkiye (Tü BI?TAK), project number 122S347 [to N.M.], European Molecular Biology Organization (EMBO), project numbers EMBO IG-5352 [to NM] and IG-4163 [to OA].

## Notes

https://brendo.sabanciuniv.edu/

### Competing Interest Statement

The authors have declared no competing interest.

