## Supplementary Info for "Brendo: an open resource for uniquely transcribed genes in brain endothelial cells"

**Supplementary Figure 1: Sample relationships for the curated dataset.** Samples were clustered using single linkage based on pairwise cosine similarities of MRN normalized counts. Rows and columns represent individual samples. Bottom color bars indicate BBB-derived samples (BBB) and sample source (Source).

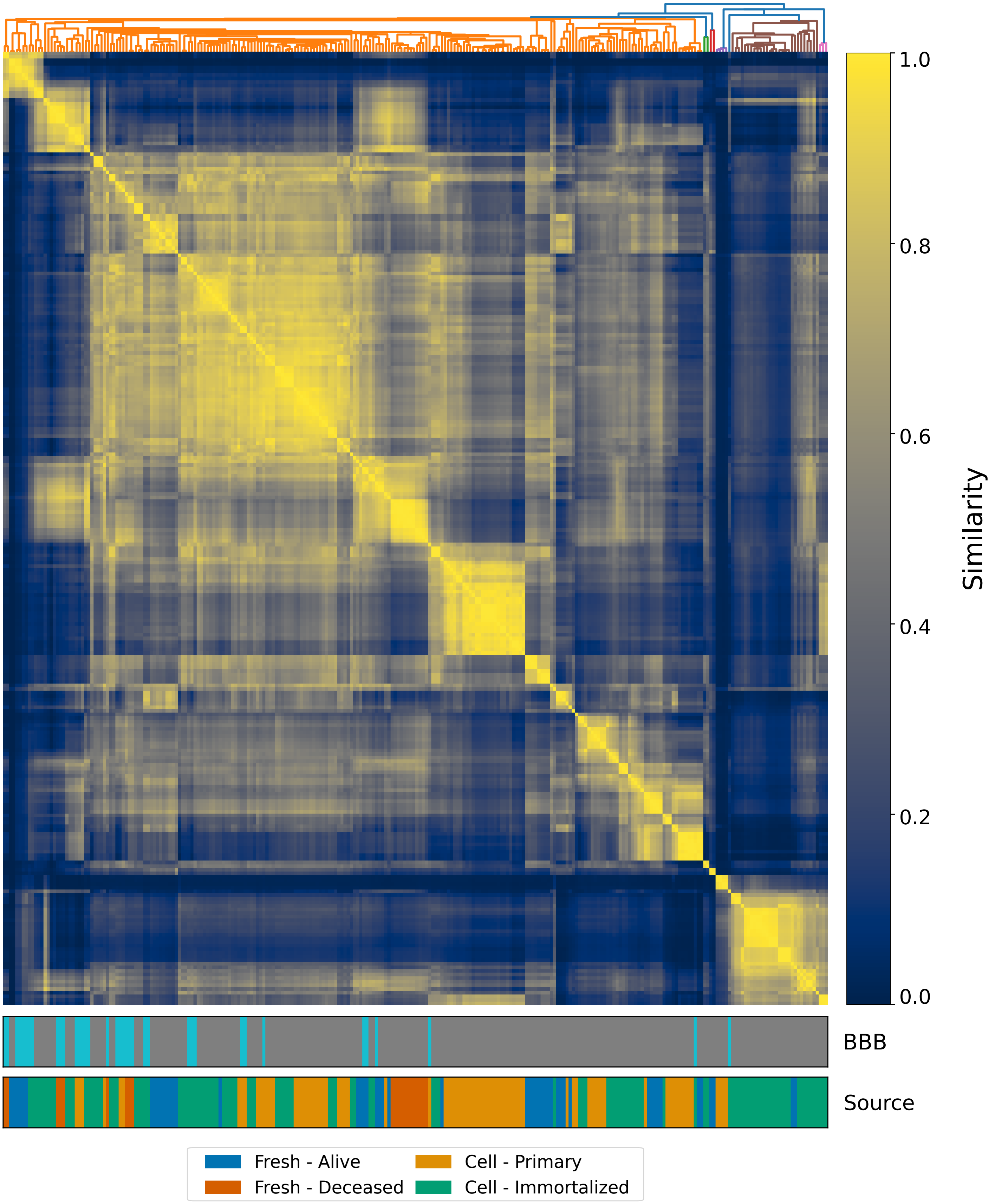

EXPORT ALL GENES EXPORT SELECTION

| Gene ID | Gene Name | Uniprot ID | Significance | DESeq2 Aggregate | DESeq2 Bootstrap | WRST Aggregate | WRST Bootstrap | Protein Evidence |
|---|---|---|---|---|---|---|---|---|
| ENSG00000173705 | SUSD5 | O60279 | **6.9987657192** | ✔ | ✔ | ✔ | ✔ | ✘ |
| ENSG00000084453 | SLCO1A2 | P46721 | **6.2385380585** | ✔ | ✔ | ✔ | ✔ | ✘ |
| ENSG00000126709 | IFI6 | P09912 | **6.0081376339** | ✔ | ✔ | ✔ | ✔ | ✘ |
| ENSG00000112414 | ADGRG6 | Q86SQ4 | **5.8354848478** | ✔ | ✔ | ✔ | ✔ | ✘ |
| ENSG00000175471 | MCTP1 | Q6DN14 | **5.2335677591** | ✘ | ✔ | ✔ | ✔ | ✘ |
| ENSG00000142235 | LMTK3 | Q96Q04 | **5.0275141321** | ✔ | ✔ | ✔ | ✔ | ✘ |
| ENSG00000133106 | EPSTI1 | Q96J88 | **4.5073913302** | ✔ | ✔ | ✔ | ✔ | ✘ |
| ENSG00000118777 | ABCG2 | Q9UNQ0 | **4.3670944841** | ✔ | ✔ | ✔ | ✔ | ✔ |
| ENSG00000070526 | ST6GALNAC1 | Q9NSC7 | **4.3009782533** | ✔ | ✔ | ✔ | ✔ | ✘ |
| ENSG00000110324 | IL10RA | Q13651 | **4.035147906** | ✔ | ✔ | ✔ | ✔ | ✘ |

Rows per page 10 1-10 of 328

**Supplementary Figure 2: Screenshot of RRA and bootstrap results from Brendo** RRA and bootstrap validation results are listed for top 10 transmembrane helix filtered genes. Additional results are available at the platform webpage.

**Supplementary Table 1:** Metadata for all samples included initially in the dataset.

| GEO | PMID | Tissue |
|---|---|---|
| GSE149699 | 35258704 | Arterial ECs |
| GSE124402 | 30852379 | Cardiac ECs |
| GSE168534 | 34480898 | HUVECs |
| GSE108919 | NA | Glomerular / Peritubular ECs |
| GSE176481 | NA | Corneal ECs |
| GSE102498 | 31539914 | Arterial ECs |
| GSE164829 | 34030460 | Pulmonary Arterial ECs |
| GSE89475 | 30651639 | Arterial ECs |
| GSE153694 | 33921957 | HUVECs |
| GSE107574 | 29234067 | Endoneurial ECs |
| GSE121444 | 32402127 | Dermal ECs, BMECs |
| GSE90489 | 28654985; 31194824 | Corneal ECs |
| GSE174545 | NA | Dermal ECs |
| GSE93705 | 29170757 | Cardiac ECs |
| GSE161021 | 33948992 | Arterial ECs |
| GSE85782 | 27804979 | Arterial ECs, HUVECs |
| GSE172210 | NA | HUVECs |
| GSE114607 | 30240741 | Multiple* |
| GSE160611 | 33468662 | Arterial ECs |
| GSE93143 | 29074968 | BMECs |
| GSE164396 | 37846557 | HUVECs |
| GSE78540 | 22955616 | Pulmonary Arterial ECs |
| GSE151312 | NA | Cardiac ECs |
| GSE176479 | 34516880 | Cardiac ECs |
| GSE131047 | 32970335 | ECs from foreskin tissue |
| GSE154546 | 33106502 | HUVECs |
| GSE155449 | 32946121 | HUVECs |
| GSE136717 | 32071273 | ECs from foreskin tissue |
| GSE140898 | 32896271 | Arterial ECs |
| GSE125907 | 34402643 | BMECs |
| GSE120732 | 33087715 | HUVECs, Peripheral Blood ECs |
| GSE93330 | 28298524 | Pulmonary Arterial ECs |
| GSE142209 | 32704093 | BMECs |
| GSE163827 | 33523764 | BMECs, Pulmonary Arterial ECs |
| GSE115954 | 31976858 | HUVECs |
| GSE138734 | 34140680 | Colonic MECs, Tonsil ECs |
| GSE143148 | 32034099 | HUVECs |
| GSE93904 | 30889293 | Mucosal MECs |
| GSE138113 | 32075753 | BMECs |
| GSE183631 | 34665379 | Dermal ECs |
| GSE138114 | 32075753 | BMECs |
| GSE185479 | 34282213 | Pulmonary Arterial ECs |
| GSE115536 | 30983154 | HUVECs |
| GSE164799 | 34072510 | Liver Sinusoidal ECs |
| GSE138628 | 31712416 | HUVECs |
| GSE179923 | 35063125 | BMECs |
| GSE137786 | 31836710 | HUVECs, Glomerular / Peritubular ECs |
| GSE145232 | 32697191 | Pulmonary Arterial ECs |
| GSE124494 | 31402260 | Lymph Node LECs |
| GSE143353 | 32238801 | Arterial ECs |
| GSE97575 | 29134197 | BMECs |
| GSE139110 | NA | Liver Sinusoidal ECs, HUVECs |
| GSE100242 | 28842720 | HUVECs |
| GSE99914 | NA | Glomerular / Peritubular ECs |
| GSE94064 | 28188910 | BMECs |
| GSE130727 | 31251810 | Arterial ECs, HUVECs |
| GSE54912 | 24732587; 28886276; 29657262 | Arterial ECs |
| GSE129933 | 31156626 | Liver Sinusoidal ECs |
| GSE167959 | 33542154 | BMECs |
| GSE125742 | 30903824 | HUVECs |
| GSE133751 | 32232013 | HUVECs |
| GSE89787 | 30582451 | Pulmonary Arterial ECs |

* : Glomerular / Peritubular ECs, Liver Sinusoidal ECs, Pulmonary Arterial ECs, Cardiac ECs.
